# Intrauterine position probabilities in mice, rats and gerbils

**DOI:** 10.1101/173153

**Authors:** John F Mulley, Ludmila I Kuncheva

## Abstract

The position of a developing embryo or foetus relative to members of the same or opposite sex can have profound effects on its resulting anatomy, physiology and behavior. Here we treat intrauterine position as a combinatorial problem and determine the theoretical probability of having 0, 1 or 2 adjacent foetuses of the opposite sex for species with random and biased distribution of genders in uterine horns (mice and gerbils), and where the influence of an “upstream” male has been proposed to be a factor (rats). As overall litter size increases the probabilities of having 0, 1, or 2 adjacent foetuses of the opposite sex approaches and eventually settles at 0.25, 0.5, 0.25 respectively. However, at biologically-relevant litter sizes probabilities are more variable and the general effect of an increase in litter size is to increase the probability that any particular foetus will be flanked by two members of the opposite sex. When gender ratios within a uterine horn are no longer balanced, the probability that there are 0 adjacent foetuses of the opposite sex increases.

## INTRODUCTION

The position of a developing embryo or foetus relative to others of the same or opposite sex can influence subsequent anatomy, physiology and behaviour. Such effects have been recognised since antiquity, even from animals that produce few offspring, such as cows, where intersex “freemartins” are formed when the placentae of male and female twins establish vascular connections enabling exchange of cellular material, resulting in chimerism and masculinisation of the reproductive tract of the female (Freeman, 2007; Hunter, 1779; Padula, 2005). In animals which produce large numbers of offspring at one time, the effects can be profound or nuanced, depending on the position of an embryo relative to members of the same or opposite sex, and are not the result of chimerism. This intrauterine position effect has been particularly well studied in rodents, and is caused by the transfer of readily diffusible lipophilic steroid hormones such as testosterone and estradiol between adjacent embryos (Clemens, 1974; Clemens et al., 1978; Nagel and vom Saal, 2004; Ryan and Vandenbergh, 2002; Vandenbergh, 2009; vom Saal, 1989; vom Saal et al., 1999, 1983; vom Saal and Bronson, 1980). Embryos located centrally within a horn of the bicornuate rodent uterus may have 0, 1 or 2 adjacent embryos of the opposite sex, and those in terminal positions (located adjacent to the cervix or ovary) 0 or 1 (Figure 1). Embryos may therefore be defined as: 2F when they develop between two females (equivalent to 0M); 2M when they develop between two males (equivalent to 0F); or 1M/1F when the develop between one neighbour of each sex. Terminal embryos can only be 1M (i.e. 0F) or 1F (0M). The most extreme phenotypic effects are seen for those that develop between two members of the opposite sex (2F males and 2M females), as these receive the greatest supplement of additional hormone (Nagel and vom Saal, 2004). 2M females exhibit an extended anogenital region, delayed puberty, decreased attractiveness, irregular oestrus cycles and decreased reproductive capacity, and increased aggression compared to 2F females, which have more regular and longer oestrus cycles, are more sexually attractive and receptive to males, experience earlier puberty, and produce more litters over their lifetime (Clark and Galef, 1998; McDermott et al., 1978; Nagel and vom Saal, 2004; Quadagno et al 1987; Ryan and Vandenbergh, 2002; Vandenbergh, 2009; Vandenbergh and Huggett, 1995; vom Saal et al., 1983; vom Saal, 1989; vom Saal et al., 1999; vom Saal and Bronson, 1980). Furthermore, the intrauterine position of a female can influence the sex ratio of her own litters, with 2M females producing litters biased towards males, and 2F females producing more female offspring (Clark et al., 1993; Vandenbergh, 1993; Vandenbergh and Huggett, 1994). Finally, male embryos that develop between two males (2M/0F males) and those that develop between two females (2F/0M) can be considered to be “studs” and “duds” respectively, with studs both more attractive to females and more reproductively successful than duds (Clark et al., 1992). Imaginary versions of real-world situations are often used as the basis of mathematical problems, especially in the field of combinatorial analysis (combinatorics). For example, the ménage problem (problème des ménages) considers the number of possible ways to seat *n* male-female couples at a circular table, such that men and women alternate, and no-one sits next to their partner (Bogart and Doyle, 1986; Dutka, 1986). The arrangement of embryos within uterine horns of litter-bearing species is a real-world situation that can also be considered as a combinatorial problem (an embryonic dinner party?).

**Figure 1.**
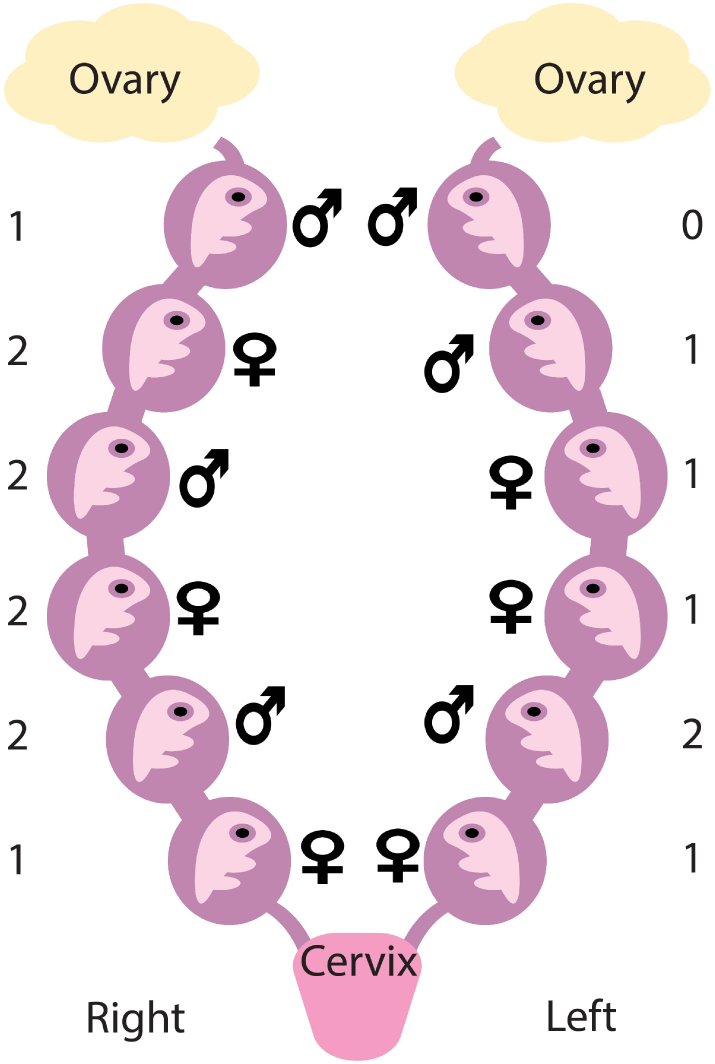
Ventral view of a bicornuate reproductive system, showing the left and right uterine horns. In this example, there are 12 foetuses (6 female ♀, 6 male ♂, divided equally between the two horns, and randomly distributed within them). Those in terminal positions (neighbouring the cervix or ovary) may have 0 or 1 adjacent foetuses of the opposite sex, and those in central positions may have 0, 1, or 2. The numbering at the far left and right of the figure highlights the situation for these foetuses.

Whilst much of the above holds true for all rodents, there are subtle differences in specific lineages, such as laboratory rats (*Rattus norvegicus*) and Mongolian gerbils (*Meriones unguiculatus*). In rats, both venous and arterial blood flows unidirectionally from the caudal end towards the cephalic end of each uterine horn (i.e. cervix to ovary), and so, in addition to effects resulting from testosterone produced by immediate neighbours, testosterone produced by male foetuses at the caudal end has been suggested to influence “downstream” littermates (Hernandez-Tristan et al., 1999; Meisel and Ward, 1981) (although it was subsequently shown that hormones move in both directions (Even et al., 1992; Vom Saal and Dhar, 1992). In gerbils, there is a biased distribution of male and female foetuses in the uterine horns, with males more common in the right horn than the left (Clark and Galef, 1990). There are clearly several factors that influence the probability that an embryo will have 0, 1 or 2 neighbours of the opposite sex, including overall litter size, the sex ratio of the litter, and the distribution of foetuses in the two uterine horns. In rats, we must also consider the presence of at least one male downstream, as a single male located caudally can have the same effect as two or more (Meisel and Ward, 1981). Here we use probability arguments and combinatorial analysis to determine probabilities of embryos being adjacent to 0,1 or 2 members of the opposite sex for random (mouse) and biased (gerbil) distribution of embryos in uterine horns, and for the “upstream male” effect for a variety of biologically-relevant (and impossible) litter sizes.

## RESULTS

### Random and fixed gender approaches

We developed two models for determining the probability that any given foetus has 0,1 or 2 adjacent members of the opposite sex.

### Case 1: Random gender

In this first model, which we call *Case 1: Random gender*, we assume that there is a fixed probability (*p*) that a randomly picked foetus is female. The exact ratio of male:female foetuses in the horn of the uterus is not fixed. For example, according to this assumption, if there are a total of 6 foetuses in the horn, and the probability of being female is 0.5, they can all be male or all can be female with a small but non-zero probability. The total number of foetuses in the horn is denoted by *K*. For this model, the probabilities for 0, 1 or 2 neighbours of the opposite sex are:

> 0 adjacent foetuses of the opposite sex: 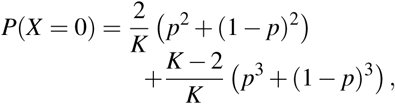
>
> 1 adjacent foetuses of the opposite sex: 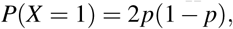
>
> 2 adjacent foetuses of the opposite sex: 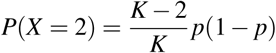

The derivation is detailed in the Methods section, Figure 2 illustrates the result for a selection of values of *K*, with balanced sex ratios (i.e. *p*=0.5), and tabulated results are provided in the Supplemental material for *K* varying from 2 to 24, and *p* ϵ {0.5,0.45,0.3}. MATLAB code and an Excel probability calculator are available at on GitHub (DOI:10.5281/zenodo.838435,https://github.com/JFMulley/Intrauterine_position). Figure 3 shows the probabilities for 0, 1 and 2 neighbours of the opposite sex as functions of the number of foetuses *K*. The sum of the three probabilities for each value of *K* is 1.

**Figure 2.**
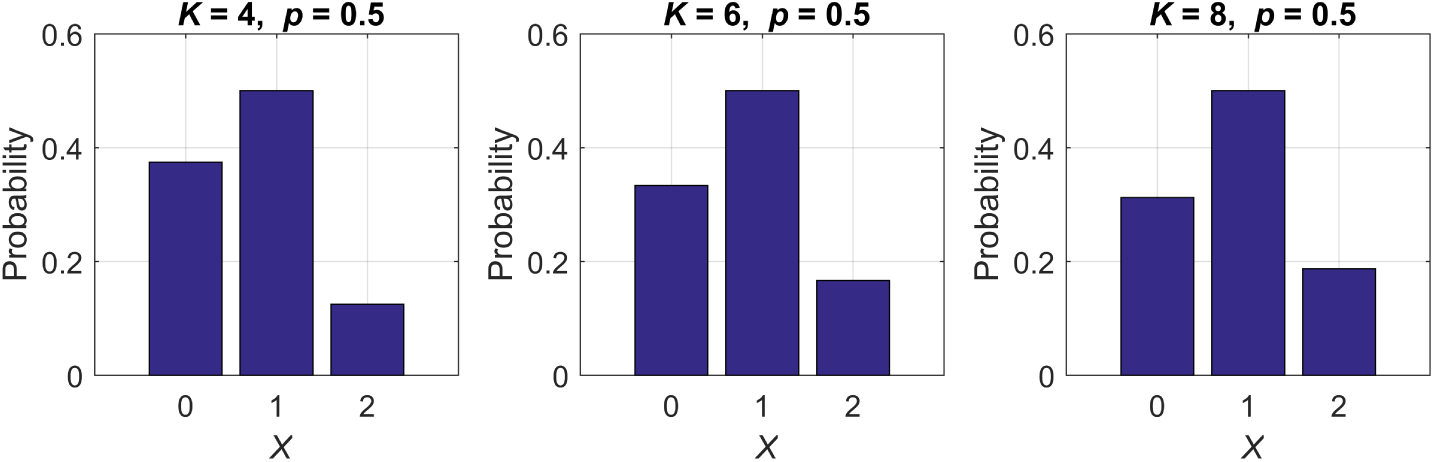
Example probability calculations for *Case 1: Random gender*. *X* is the number of adjacent foetuses of opposite sex, *K* is the number or foetuses in the uterine horn, and *p* is the probability that a randomly chosen foetus is female (here set at *p*=0.5 to reflect a balanced sex ratio).

**Figure 3.**
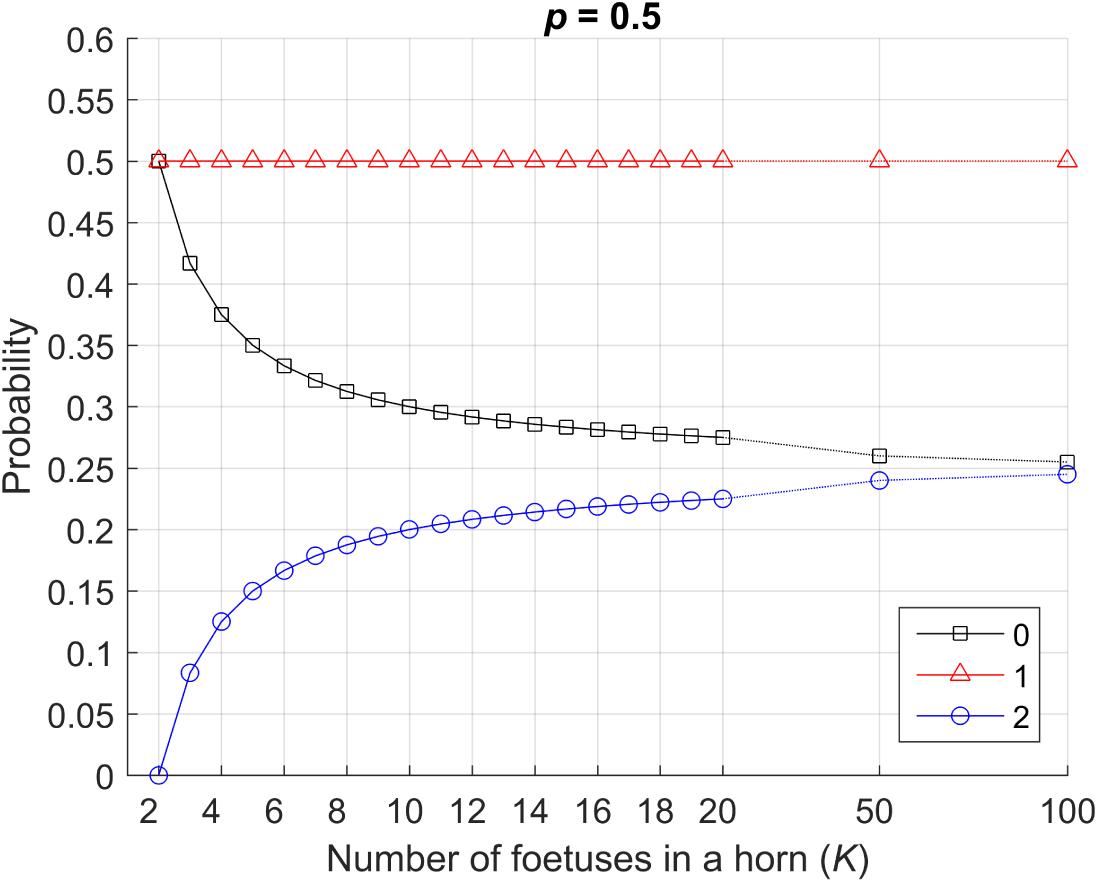
Probability of 0, 1 and 2 adjacent foetuses of the opposite sex as a function of the number of foetuses in the uterine horn for *Case 1: Random gender*, with the probability (*p*) that a randomly picked foetus is female set at 0.5 (i.e. a balcnced sex ratio). Increasing litter size has no impact on the probability that there is at least one adjacent foetus of the opposite sex, but does increase the probability that a foetus will be flanked by two members of the opposite sex. The asymptotic values are 0.25,0.5,0.25 for 0, 1, and 2 adjacent foetuses of the opposite sex, respectively.

The probability that there is 1 neighbour of the opposite sex does not depend on the number of foetuses *K*, and is always stable at 0.5. The other two probabilities converge asymptotically to the following values:

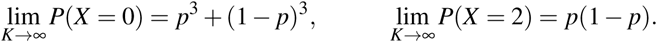

For *p* = 0.5 (equal numbers of males and females), these limits are 0.25,0.5,0.25 for 0, 1, and 2 adjacent foetuses of the opposite sex, respectively.

### Case 2: Fixed gender

For the second model, *Case 2: Fixed gender*, we assume that there is a fixed number of female foetuses (*n*) among the *K* foetuses in the uterine horn. The probabilities for 0, 1 or 2 adjacent foetuses of the opposite sex for this model are:

> 0 adjacent foetuses of the opposite sex: *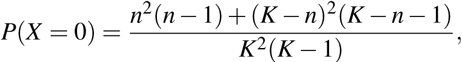*
>
> 1 adjacent foetuses of the opposite sex: *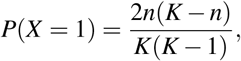*
>
> 2 adjacent foetuses of the opposite sex 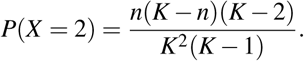

The derivation is detailed in the Methods section as before, Figure 4 depicts similar examples to those in Figure 2, and tabulated results are given in the supplementary material.

**Figure 4.**
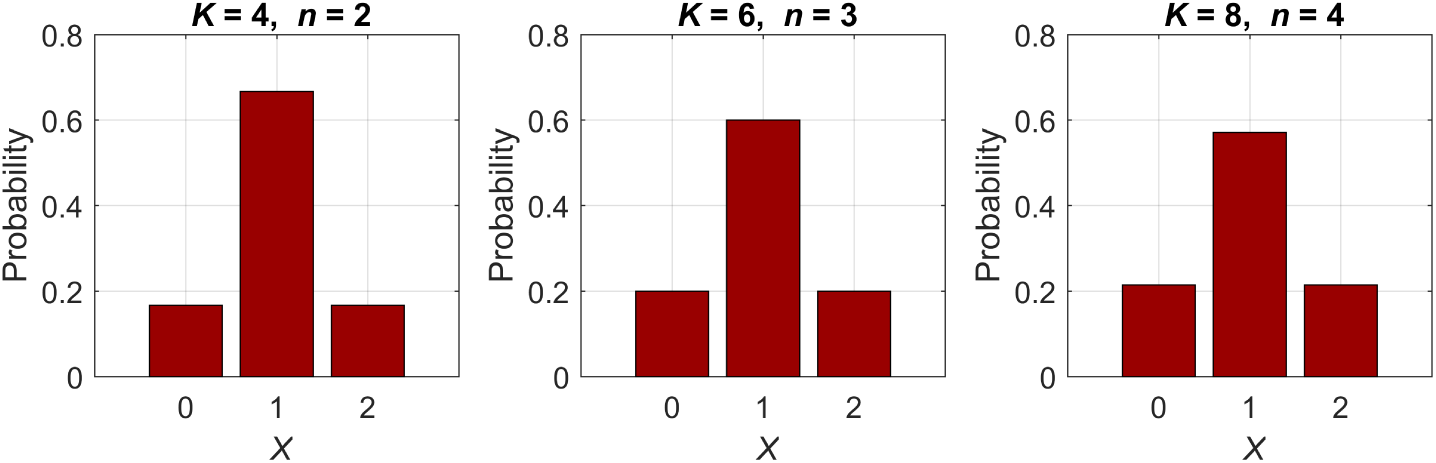
Examples of probability calculations for *Case 2: Fixed gender*. *X* is the number of adjacent foetuses of the opposite sex, *K* is the number of foetuses in the uterine horn, and *n* is the number of female foetuses (here set at *K*/2 to reflect a balanced sex ratio in each uterine horn).

The probability of 0, 1 and 2 adjacent foetuses of the opposite sex as a function of the number of foetuses in the uterine horn for *Case 2: Fixed gender* is shown in Figure 5. The number of females was taken to be *K/*2 (*exactly* half, as in a litter with a balanced sex ratio in each uterine horn). There is a pronounced difference between the curves in Figures 3 and 5. This can be explained with the different assumptions. Take, for example, *K* = 2 foetuses, *p* = 0.5 and *n* = *K/*2. In Case 1, the probability of having a neighbour of the opposite sex is exactly 0.5. In case 2, however, we *know* that one of the foetuses is male, and the other is female. Then the probability of having a neighbour of the opposite sex is 1. The dramatic differences between the probability curves highlights the importance of specifying the assumptions and the model when quoting probabilities in this context. Asymptotically (K → ∞) both Case 1 and Case 2 converge to the same limit values: *P*(*X* = 0) = 0.25, *P*(*X* = 1) = 0.25, and *P*(*X* = 2) = 0.25.

**Figure 5.**
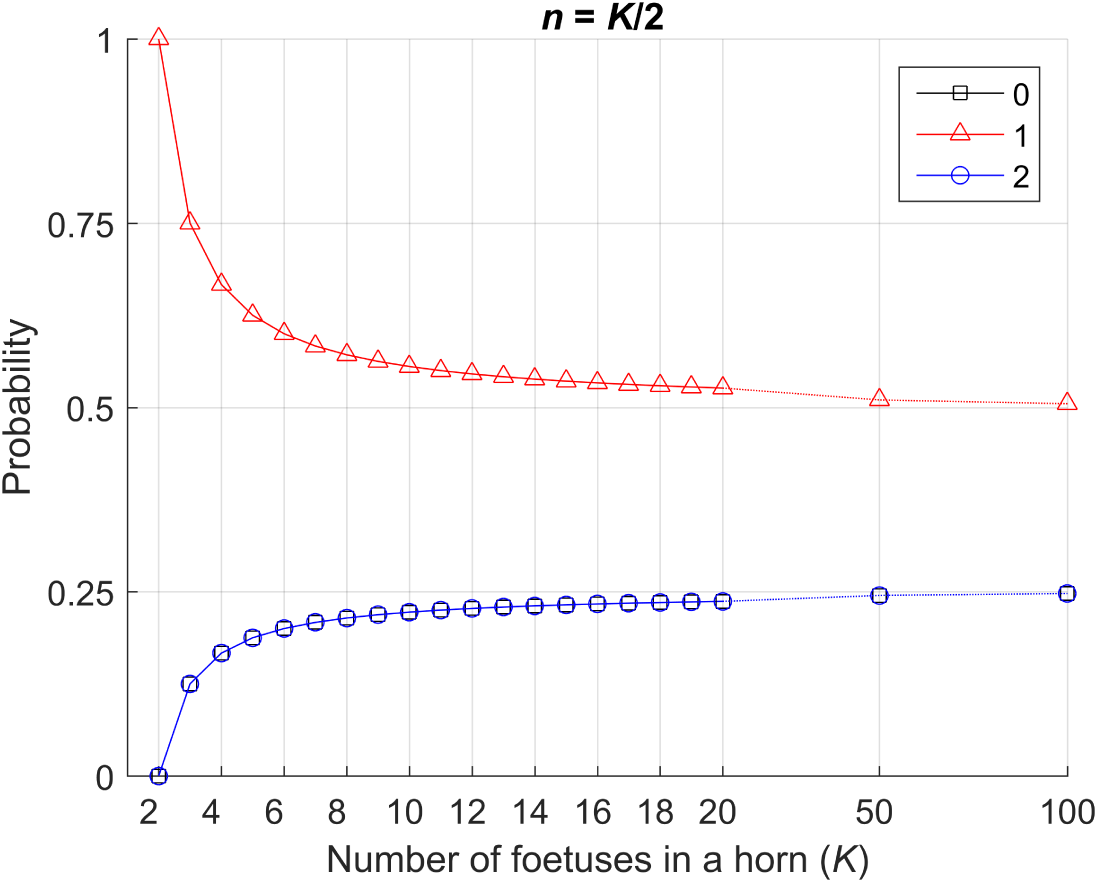
Probability of 0, 1 and 2 adjacent foetuses of the opposite sex as a function of the number of foetuses in the uterine horn for *Case 2: Fixed gender*. Here the number of female foetuses *n* is set at *K*/2 to reflect a balanced sex ratio. The asymptotic values are 0.25,0.5,0.25 for 0, 1, and 2 adjacent foetuses of the opposite sex, respectively. The curves for 0 and 2 coincide.

### Changing the gender ratio

Where gender ratios within the uterine horn are equal, the probabilities of a given foetus being located adjacent to 0, 1 or 2 members of the opposite sex settle at 0.25, 0.5 and 0.25 respectively (Figures 3 and5). When gender ratios are no longer balanced (as in the gerbil, where male embryos are more common in the right horn (Clark and Galef, 1990)), the probability of being adjacent to a member of the opposite sex decreases, and the probability of being adjacent to 0 members of the opposite sex increases (Figure 6). For Case 1, if *p* = 0.3, then the probabilities for 0, 1, and 2 neighbours of the opposite sex would be 0.37,0.42,0.21, respectively.

**Figure 6.**
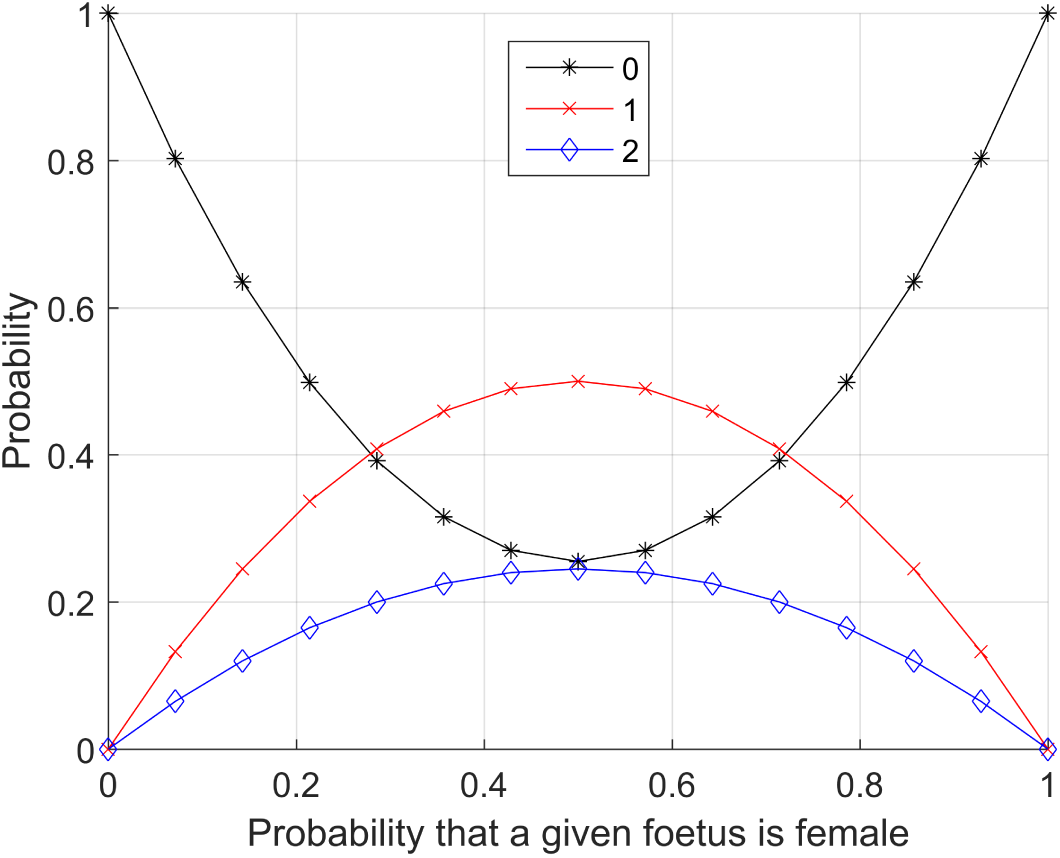
Probability of 0, 1 and 2 adjacent foetuses of the opposite sex as a function of the probability of female, *p*, calculated from *Case 1: Random gender*. At *p*=0.5 the sex ratio is equal and the probabilities that a given foetus is adjacent to 0, 1 or 2 members of the opposite sex are 0.25, 0.5 and 0.25 respectively. As sex ratios within the uterine horn become biased towards one sex or the other (i.e. shifts left or right on the x axis), the probability of being adjacent to 1 or 2 members of the opposite sex decreases (and therefore the probability of being adjacent to a member of the same sex increases).

A similar calculation can be performed for Case 2, where the number of foetuses in a uterine horn *K* increases but the number of female foetuses *n* is fixed (e.g. *n*=3, Figure 7). The graph starts at *K*=3 because there are necessarily *n* = 3 females. At *K* = 6, the probabilities are 0.2, 0.6 and 0.2, and from *K* = 3 to *K* = 6, the probability of having neighbours of the same sex (*X* = 0, black dashed line) decreases because 3 male foetuses are gradually introduced. From this point onwards, however, the graph goes upwards because the population is dominated by males, and the male proportion increases with each increment of *K*. Consequently, the females will become progressively rarer, and the probability of 1 or 2 female neighbours will decrease with increasing *K*. This example illustrates the asymptotic case where K → ∞, and *n* stays a constant. The probabilities for these asymptotic cases are respectively *P*(*X* = 0) = 1, *P*(*X* = 1) = 0, and *P*(*X* = 2) = 0.

**Figure 7.**
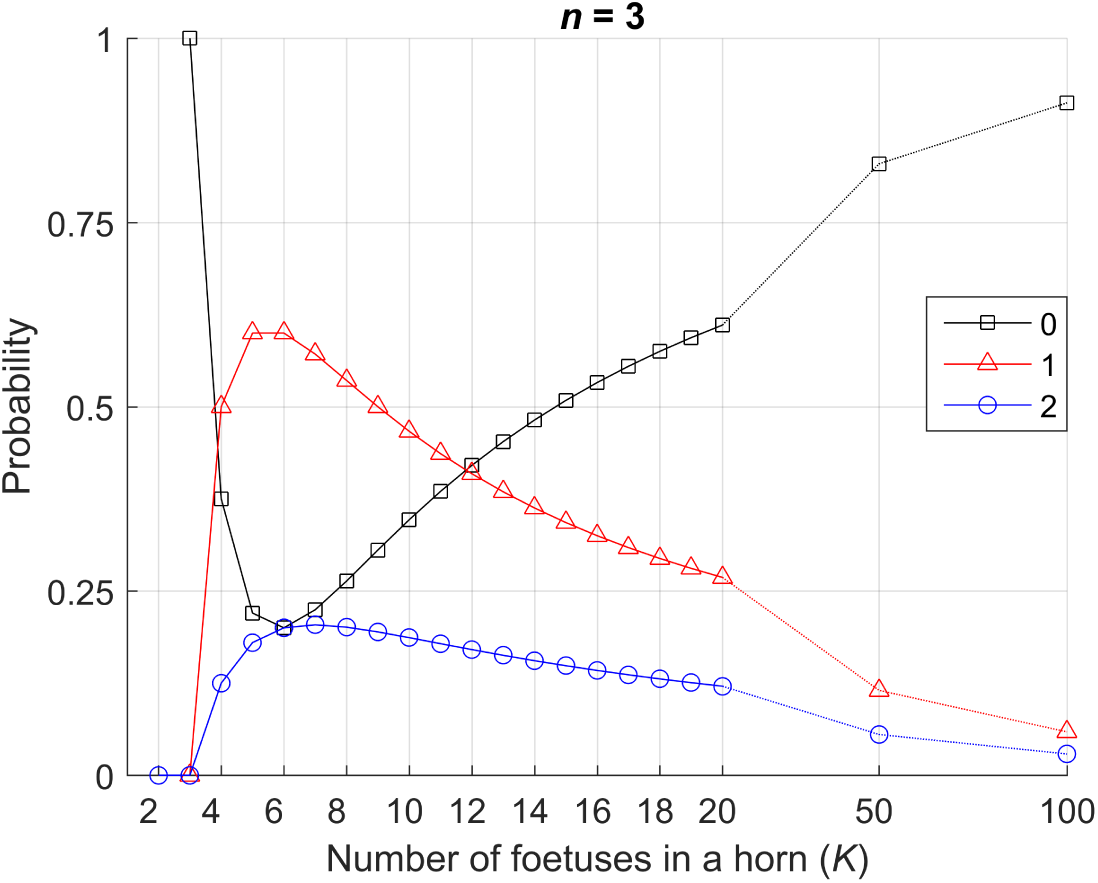
Probability of 0, 1 and 2 adjacent foetuses of the opposite sex as a function of the number of foetuses in the uterine horn for *Case 2: Fixed gender* for a constant number of females *n* = 3. When *K* and *n* are equal, all foetuses are female and there is no possibility that an adjacent foetus is a member of the opposite sex. From *K* = 3 to *K* = 6, the probability of having neighbours of the same sex decreases. At *K*=6 the sex ratio within the uterine horn is balanced, but since *n* is fixed, all subsequent additions are male, and at values of *K >*6 a male bias develops.

### ”Upstream males”

The unidirectional (cervix to ovary) flow of blood in the rat uterus adds a further complication, as the testosterone produced by a single male was suggested to be sufficient to inuence all downstream embryos (Hernandez-Tristan et al., 1999; Meisel and Ward, 1981). Whilst it was subsequently shown that hormone movement is bidirectional (Even et al., 1992; Vom Saal and Dhar, 1992), the “upstream male” hypothesis represents an interetsing thought expeirment, and so we therefore next developed an approach to account for this. In a uterine horn containing *K* foetuses, ordered 1 – 2 – 3 – 4 – … – *K*, and a foetus at position *y*, we visualise the probability that there is at least one male at any position between 1 and *y* – 1 (the argument is detailed in the Methods section). Figure 8 shows the probability of at least one “upstream” male as a function of the position of the foetus of interest *y*, and *K* = 12, for both models. For *Case 1: Random gender*, in a balanced (*p*=0.5) sex ratio, the probability of an upstream male raises to 99% for *y ≥* 8, and to above 99.9% for the last position. For *Case 2: Fixed gender*, an upstream male is a certainty earlier, as soon as *y* reaches *n* + 1, where *n* is the number of females in the uterine horn (Figure 8).

**Figure 8.**
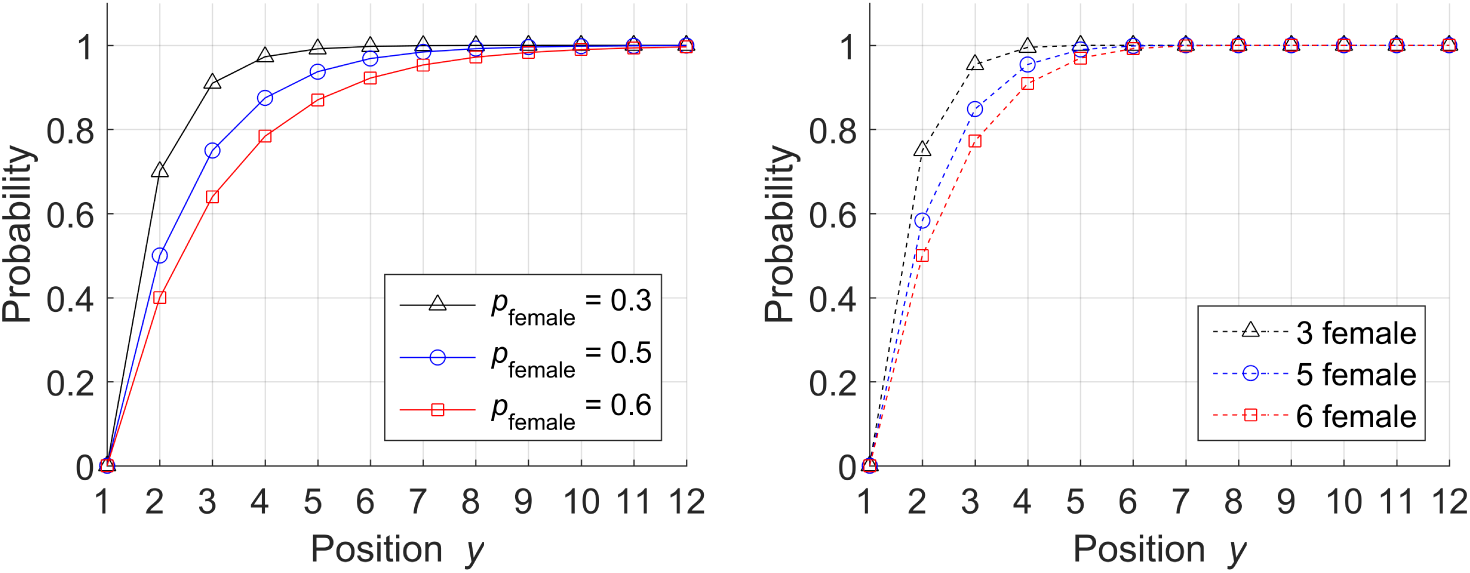
Probability of at least one “upstream male”. for *Case 1: Random gender* (left), and *Case 2: Fixed gender* (right) for *K* = 12 foetuses in a horn. *y* is the position of the foetus of interest.

## DISCUSSION

After decades of neglect, the importance of sex as a biological variable (SABV) in basic, preclinical and clinical research is gaining increasing recognition (Clayton and Collins, 2014; Cornelison and Clayton, 2017; Johnson et al., 2009; Miller et al., 2017). For studies using rodents, it will also be important to consider the impact of intrauterine position on embryonic development and adult morphology and behavior, and to facilitate this we have provided data, tools and resources for the determination of intrauterine position probabilities for various litter sizes and sex ratios in several common model species. Our results and approaches will also be applicable other rodent species, such as spiny mice (*Acomys sp*. a model for tissue regeneration (Santos et al., 2016; Seifert et al., 2012)), deer mice (*Peromyscus sp.*, models for population genetics and adaptation (Weber et al., 2013; Bedford and Hoekstra, 2015; Bendesky et al., 2017)), sandrats (a model for diet-induced diabetes (Hargreaves et al., 2017; Donath et al., 1999)), hamsters (Brekke et al., 2016; Brekke and Good, 2014) and degus (*Octodon degus* (Roff et al., 2017; Correa et al., 2016, 2013)), as well as rabbits (Banszegi et al., 2009) and other mammals.

It has been known for several decades that IUP has significant effects on reproductive success, and on sex ratios in subsequent generations (Clemens et al., 1978; Vandenbergh, 2009; Ryan and Vandenbergh, 2002). We should therefore be extremely careful when using generalisations regarding “males” and “females”, as clearly not all members of these categories are equal. Such considerations may be useful in the field of sex allocation and sex ratio theory, where much effort has been expended on the evolutionary and adaptive significance of alterations from a 1:1 ratio of males and females, especially with respect to sociality and altruism (West et al., 2000, 2002; West and Sheldon, 2002; Wild and West, 2007; Charnov, 1981; Fisher, 1930). In social species such as gerbils, IUP may be the determining factor in assigning roles at the nest, with asexual 2F “dud” males equivalent to a cadre of sterile helpers (Clark and Galef, 2000; Downing et al., 2017) and aggressive, highly sexual 2M “stud” males important for securing territory, and for dispersal. 2F females reach puberty at an earlier age, are more attractive to males, and produce more litters over their lifetime than 2M females (vom Saal et al., 1999; Ryan and Vandenbergh, 2002; Nagel and vom Saal, 2004). The biased distribution of males and female foetuses in the uterine horns of gerbils results in a decrease in the the number of foetuses that develop adjacent to a member of the opposite sex (Figure 6 and Figure 7), and so will produce a greater proportion of stud (2M) males and “super-mother” (2F) females. Whilst some probability values have been reported in the literature previously (e.g. (Nagel and vom Saal, 2004; vom Saal, 1981; Clark and Galef, 1990; vom Saal and Bronson, 1980; vom Saal, 1989; Clark et al., 1991)), to our knowledge there have been no previous attempts to quantify the probability that there is at least one upstream male (Figure 8). For the IUP probabilities reported previously, the approach and underlying assumptions used to generate these values have not been provided. As we have shown, different assumptions can generate quite different results - for example, when there are six foetuses in a uterine horn and three of these *must* be female (as in our *Case 2: Fixed gender*), the probabilities of any foetus having 0, 1, or 2 adjacent foetuses of the opposite sex are 0.2, 0.6 and 0.2 respectively (Figure 4 and Supplemental Table 2). However, when there are 6 foetuses in the horn and the probability that any one of these is female =0.5 (i.e. theoretically, the gender ratio within the uterine horn is balanced), the values are 0.333, 0.5 and 0.1667 (Figure 2 and Supplemental Table 1). The situation is further complicated by seemingly impossible values that have been claimed, such as the 1/6, 3/6 and 1/6 for 2M, 1M and 0M respectively in a litter of 12 pups reported by (Nagel and vom Saal, 2004), and the 0.2, 0.65, 0.15 values given by (Hotchkiss and Vandenbergh, 2005), which cites (vom Saal, 1981) as a source. However, the actual values given in (vom Saal, 1981) equate to our asymptotic values of 0.25, 0.5 and 0.25. We have been unable to replicate either of these sets of values under any conditions. Hopefully then, our detailed analysis under various assumptions will go some way to providing clarity in the field, and facilitate the incorporation of intrauterine position as a biological variable in scientific research.

**Table 1.**
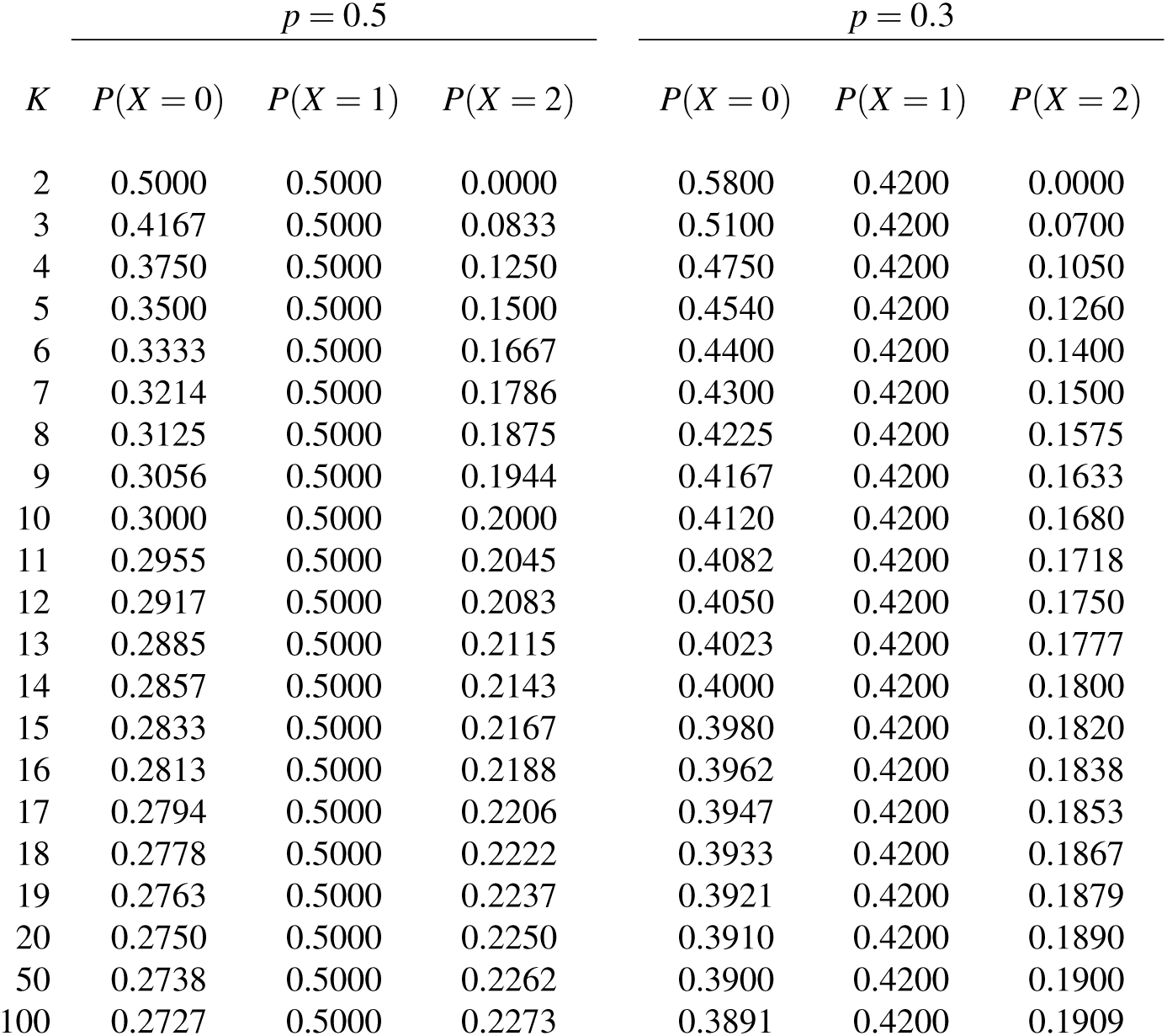
Probabilities of 0, 1 or 2 neighbours of the opposite sex for *Case 1: Random Gender*. *p* is the probability that a randomly picked foetus is female and *K* is the number of foetuses in the horn of the uterus. On the left, *p*=0.5, reflecting a balanced sex ratio. On the right, *p*=0.3, reflecting a bias towards males.

| $K$ | $p = 0.5$ | | | $p = 0.3$ | | |
| --- | --- | --- | --- | --- | --- | --- |
| | $P(X = 0)$ | $P(X = 1)$ | $P(X = 2)$ | $P(X = 0)$ | $P(X = 1)$ | $P(X = 2)$ |
| 2 | 0.5000 | 0.5000 | 0.0000 | 0.5800 | 0.4200 | 0.0000 |
| 3 | 0.4167 | 0.5000 | 0.0833 | 0.5100 | 0.4200 | 0.0700 |
| 4 | 0.3750 | 0.5000 | 0.1250 | 0.4750 | 0.4200 | 0.1050 |
| 5 | 0.3500 | 0.5000 | 0.1500 | 0.4540 | 0.4200 | 0.1260 |
| 6 | 0.3333 | 0.5000 | 0.1667 | 0.4400 | 0.4200 | 0.1400 |
| 7 | 0.3214 | 0.5000 | 0.1786 | 0.4300 | 0.4200 | 0.1500 |
| 8 | 0.3125 | 0.5000 | 0.1875 | 0.4225 | 0.4200 | 0.1575 |
| 9 | 0.3056 | 0.5000 | 0.1944 | 0.4167 | 0.4200 | 0.1633 |
| 10 | 0.3000 | 0.5000 | 0.2000 | 0.4120 | 0.4200 | 0.1680 |
| 11 | 0.2955 | 0.5000 | 0.2045 | 0.4082 | 0.4200 | 0.1718 |
| 12 | 0.2917 | 0.5000 | 0.2083 | 0.4050 | 0.4200 | 0.1750 |
| 13 | 0.2885 | 0.5000 | 0.2115 | 0.4023 | 0.4200 | 0.1777 |
| 14 | 0.2857 | 0.5000 | 0.2143 | 0.4000 | 0.4200 | 0.1800 |
| 15 | 0.2833 | 0.5000 | 0.2167 | 0.3980 | 0.4200 | 0.1820 |
| 16 | 0.2813 | 0.5000 | 0.2188 | 0.3962 | 0.4200 | 0.1838 |
| 17 | 0.2794 | 0.5000 | 0.2206 | 0.3947 | 0.4200 | 0.1853 |
| 18 | 0.2778 | 0.5000 | 0.2222 | 0.3933 | 0.4200 | 0.1867 |
| 19 | 0.2763 | 0.5000 | 0.2237 | 0.3921 | 0.4200 | 0.1879 |
| 20 | 0.2750 | 0.5000 | 0.2250 | 0.3910 | 0.4200 | 0.1890 |
| 50 | 0.2738 | 0.5000 | 0.2262 | 0.3900 | 0.4200 | 0.1900 |
| 100 | 0.2727 | 0.5000 | 0.2273 | 0.3891 | 0.4200 | 0.1909 |

**Table 2.** Probabilities of 0, 1 or 2 neighbours of the opposite sex for *Case 2: Fixed Gender*. *n* is the number of female foetuses from a total of *K* foetuses in the horn of the uterus. On the left, *n*=*K*/2 (i.e. a balanced gender ratio), and so the total number of foetuses in the horn must be even. On the right, the number of females (*n*=3) does not change as the total number of foetuses in the uterine horn increases.

| $K$ | $n = \frac{K}{2}$ | | | $n = 3$ | | |
| --- | --- | --- | --- | --- | --- | --- |
| | $P(X = 0)$ | $P(X = 1)$ | $P(X = 2)$ | $P(X = 0)$ | $P(X = 1)$ | $P(X = 2)$ |
| 2 | 0.0000 | 1.0000 | 0.0000 | — | — | — |
|  | — | — | — | 1.0000 | 0.0000 | 0.0000 |
| 4 | 0.1667 | 0.6667 | 0.1667 | 0.3750 | 0.5000 | 0.1250 |
|  | — | — | — | 0.2200 | 0.6000 | 0.1800 |
| 6 | 0.2000 | 0.6000 | 0.2000 | 0.2000 | 0.6000 | 0.2000 |
|  | — | — | — | 0.2245 | 0.5714 | 0.2041 |
| 8 | 0.2143 | 0.5714 | 0.2143 | 0.2634 | 0.5357 | 0.2009 |
|  | — | — | — | 0.3056 | 0.5000 | 0.1944 |
| 10 | 0.2222 | 0.5556 | 0.2222 | 0.3467 | 0.4667 | 0.1867 |
|  | — | — | — | 0.3851 | 0.4364 | 0.1785 |
| 12 | 0.2273 | 0.5455 | 0.2273 | 0.4205 | 0.4091 | 0.1705 |
|  | — | — | — | 0.4527 | 0.3846 | 0.1627 |
| 14 | 0.2308 | 0.5385 | 0.2308 | 0.4819 | 0.3626 | 0.1554 |
|  | — | — | — | 0.5086 | 0.3429 | 0.1486 |
| 16 | 0.2333 | 0.5333 | 0.2333 | 0.5328 | 0.3250 | 0.1422 |
|  | — | — | — | 0.5549 | 0.3088 | 0.1362 |
| 18 | 0.2353 | 0.5294 | 0.2353 | 0.5752 | 0.2941 | 0.1307 |
|  | — | — | — | 0.5937 | 0.2807 | 0.1256 |
| 20 | 0.2368 | 0.5263 | 0.2368 | 0.6108 | 0.2684 | 0.1208 |
|  | — | — | — | 0.8296 | 0.1151 | 0.0552 |
| 100 | 0.2475 | 0.5051 | 0.2475 | 0.9124 | 0.0588 | 0.0288 |

## CONCLUSIONS

The underlying assumptions regarding whether there are a fixed number of males and females (as in our *Case 2: Fixed gender*), or merely a predetermined probability of a given foetus being a particular sex (as in our *Case 1: Random gender*) can have dramatic effects on intrauterine position probabilities. However, in both cases the asymptotic values are 0.25, 0.5 and 0.25 for the probabilities that there are 0, 1 or 2 adjacent foetuses of the opposite sex. The biased distribution of embryos in the uterine horns of gerbils leads to a decrease in the probability that a foetus will be adjacent to a member of the opposite sex, and so increases the probability of same-sex neighbours.

## METHODS

This section details the derivations of the results. We will be using the following notations:

> *t*: gender of the embryo of interest, *t* ϵ {*F,M*}
>
> *v,w*: genders of the neighbours of the embryo of interest
>
> *p*: probability that a randomly chosen embryo is female *P*(*t* = *F*); hence *P*(*t* = *M*) = 1 – *p*
>
> *K*: number of embryos in the horn of the womb (arranged as 1 – 2 – 3 – 4 – … – *K*)
>
> *X*: a random variable denoting the number of neighbours of opposite sex; *X* ϵ {0,1,2}.

### Derivation of Case 1: Random gender

Here we assume that there are *K* embryos in the horn of the womb, and their gender is not known in advance. Any given embryo could be male or female, which is governed by a fixed probability *p*. The task is to find the probability mass function of *X*. Denote by *SP* the special cases where we pick embryo 1 or embryo *K*, both of whom have only one neighbour. The remaining *K* – 2 embryos have 2 neighbours each. Then

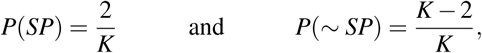

where *~* denotes the opposite event. The three values of *P*(*X*) can be calculated separately as follows:

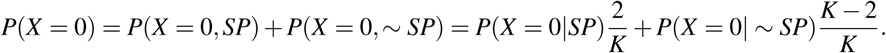

Taking into account the assumption that the genders do not depend on the position of the embryo, the conditional probabilities are respectively:

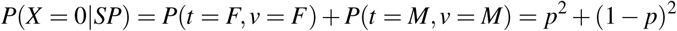

and

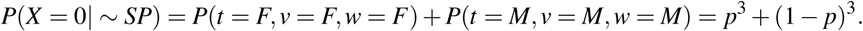

Then

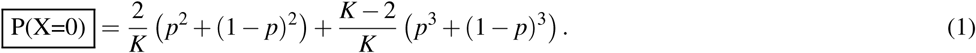

Next we can calculate *P*(*X* = 2):

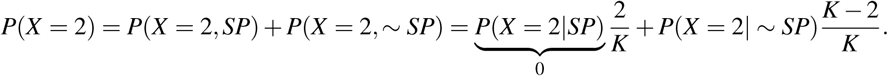

The first conditional probability is 0 because there cannot be two neighbours of different gender for the embryos at the end positions. For the other conditional probability,

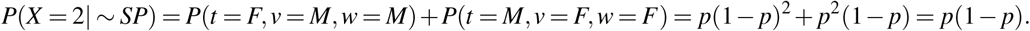

Hence,

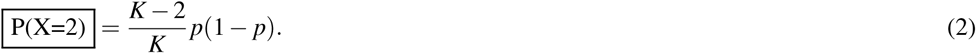

Finally,

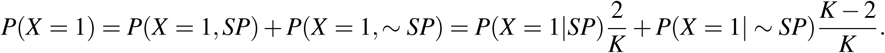

For this case,

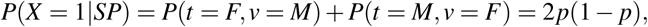

and

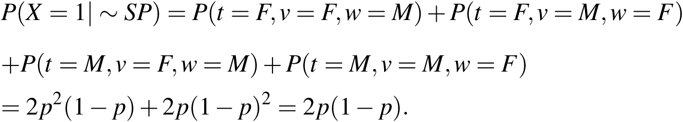

Noticing that the two conditional probabilities are identical,

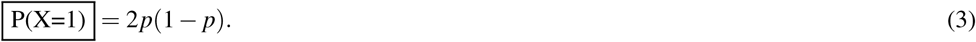

Putting the probability mass function of *X* together, the probability that the number of neighbours of different gender for a randomly chosen embryo is:

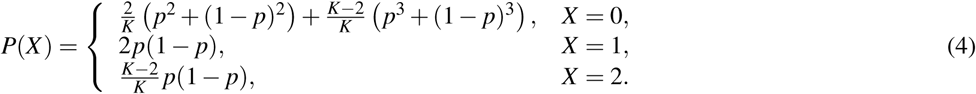

### Derivation of Case 2: Fixed gender

This time, the genders are fixed and the *K* embryos are arranged in random order. There are exactly *n* female embryos and *K – n* male embryos. The notations and the task are the same as in the previous case.

Again,

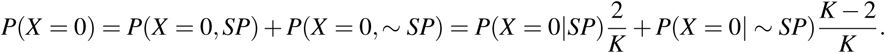

This time, however, the conditional probabilities are different:

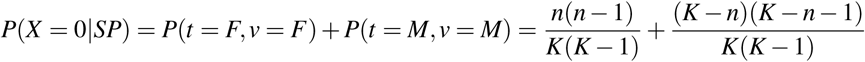

and

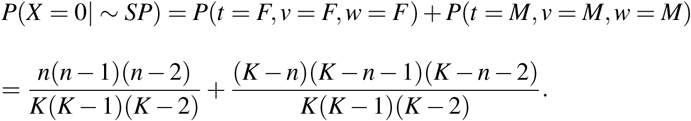

Then, after simple algebraic manipulations,

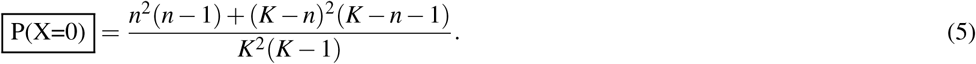

Similarly,

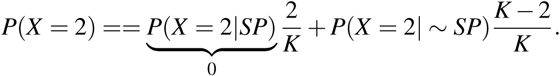

and

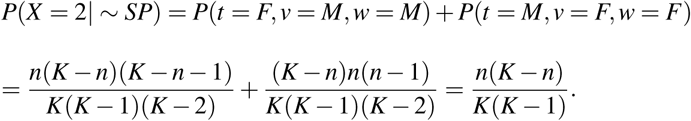

Then

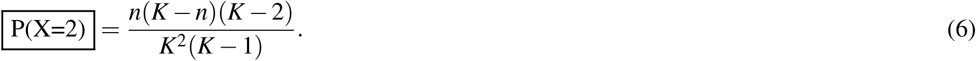

Finally,

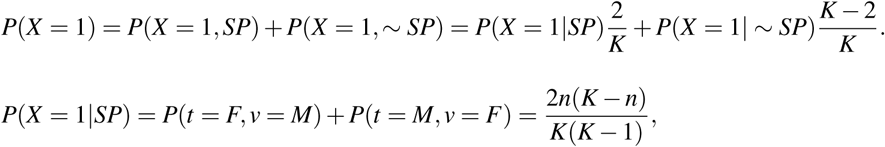

and

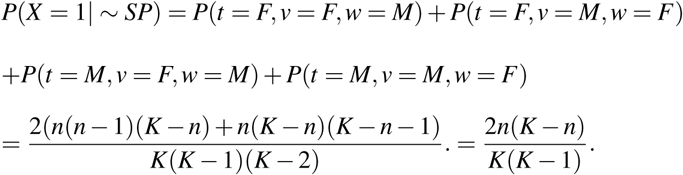

Again,

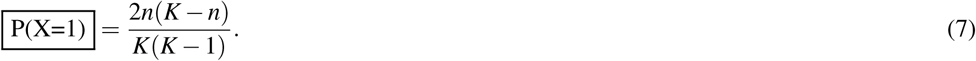

Putting equations (5), (6), and (7) together, the probability that the number of neighbours of different gender for a randomly chosen embryo is:

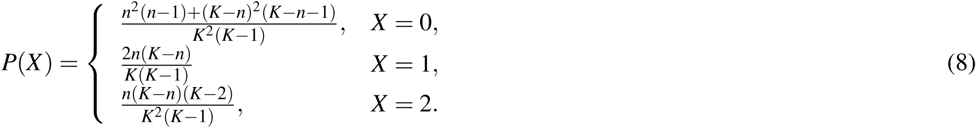

### Derivation of the probability of “upstream males”

Here we seek to answer theoretically the question: “What is the probability that an foetus at position *y* is preceded by at least one male foetus?”

In Case 1 (Random gender), the question should be phrased more precisely as: “Given the number of foetuses in the horn (*K*), and the probability of female (*p*), what is the probability that a foetus at position *y* is preceded by at least one male foetus?” Denote this probability by *PM*(*y|K, p*). *y* is a random variable taking values in the set {1,2*,…,K*}. The probability of at least one male out of *y* – 1 independent Bernoulli trials is

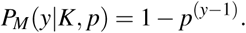

In Case 2 (Fixed gender), the question should be phrased as: “Given the number of foetuses in the horn (*K*), and the number of female foetuses (*n*), what is the probability that a foetus at position *y* is preceded by at least one male foetus?” Denote this probability by *PM*(*y|K,n*). Clearly, *PM*(*y|K,n*) = 1 for any *y > n* + 1 because there can only be *n* female foetuses among the first *y* – 1, the remaining *y* – 1 – *n* foetuses must therefore be male. When *y ≤ n* + 1, the probability of having at least one male foetus among the *y* – 1 foetuses is the opposite of the probability of all *y* – 1 foetuses being female. This probability can be calculated as the ratio of the number of combinations of *y* – 1 out of *n* and the number of all combinations of *y* – 1 out of *K*. Then the probability of at least one male out of *y* – 1 independent trials is

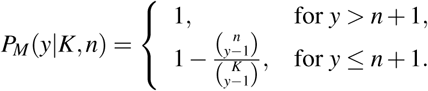

## Resources

The associated MATLAB code and an Excel format probability calculator are available on GitHub (DOI: 10.5281/zenodo.838435,https://github.com/JFMulley/Intrauterine_position). Tabulated results for various values of *p* and *n* are provided in the Supplemental material.

## ACKNOWLEDGMENTS

JFMs work on rodents is funded by a Leverhulme Trust Research Project Grant (RPG-2015-450).

